# Integrated identification and quantification error probabilities for shotgun proteomics

**DOI:** 10.1101/357285

**Authors:** Matthew The, Lukas Käll

## Abstract

Protein quantification by label-free shotgun proteomics experiments is plagued by a multitude of error sources. Typical pipelines for identifying differentially expressed proteins use intermediate filters in an attempt to control the error rate. However, they often ignore certain error sources and, moreover, regard filtered lists as completely correct in subsequent steps. These two indiscretions can easily lead to a loss of control of the false discovery rate (FDR). We propose a probabilistic graphical model, *Triqler*, that propagates error information through all steps, employing distributions in favor of point estimates, most notably for missing value imputation. The model outputs posterior probabilities for fold changes between treatment groups, highlighting uncertainty rather than hiding it. We analyzed 3 engineered datasets and achieved FDR control and high sensitivity, even for truly absent proteins. In a bladder cancer clinical dataset we discovered 35 proteins at 5% FDR, whereas the original study discovered 1 and MaxQuant/Perseus 4 proteins at this threshold. Compellingly, these 35 proteins showed enrichment for functional annotation terms, whereas the top ranked proteins reported by MaxQuant/Perseus showed no enrichment. The model executes in minutes and is freely available at https://pypi.org/project/triqler/.

## Introduction

Shotgun proteomics has in recent years made rapid advances from being a tool for large-scale identification to also include accurate quantification of proteins [23]. software packages have been developed to facilitate the quantitative interpretation of MS data, for a review see e.g. [21]. Compared to software for protein identification, the protein quantification pipelines contain many more facets, one of which actually is protein identification. Slowly but steadily the softwares for protein identification are getting their error rates under better control, though much work is still left [27, 32]. However, error rates in protein quantification have been mostly limited to setting intermediate *false discovery rate* (FDR) thresholds for the identifications or other heuristic cutoffs, such as requiring at least a certain number of peptides [8, 3] or a certain correlation between peptide quantifications [41, 40]. This gives no direct control of the errors in the reported lists of differential proteins and also discards potentially valuable information for proteins that just missed one of the thresholds. Consequently, many protein-level differential expression methods lack sensitivity, and several researchers refrain from using multiple hypothesis corrections of their summarized proteins [24]. We believe that we no longer have to accept this, as the necessary tools are already available to us in the framework of Bayesian statistics. In particular, we note that probabilistic graphical models (PGM) have the inherent ability to combine several sources of errors.

Bayesian statistics has already been used in several applications within proteomics. Most notably, it is currently being used for PSM-level identification FDR estimates [7, 12], and protein inference [29]. More recently, Bayesian methods have been applied to labeled protein quantification [28, 22]. Each of these methods has applied Bayesian statistics to parts of the quantification pipeline, but an integrated model for protein quantification is still lacking.

The need for an integrated model becomes clear once one considers the most natural hypothesis [33] for protein quantification: one strives to estimate the combined probability that a particular protein is (i) correctly identified, (ii) correctly quantified and (iii) present in a different quantity between treatment groups. The separate probabilities of (i), (ii) and (iii) are less interesting individually and worse, one is easily lulled into a false sense of reliability by claims of control of the FDR in individual steps. What we generally fail to acknowledge is that the intermediate lists are considered as fully correct by subsequent steps. The most striking example is the widely used approach of applying say a 5% protein-level identification FDR threshold, followed by a 5% differential-expression FDR threshold. Say that the protein-level identification threshold results in a list of 1000 proteins and the subsequent application of the differential-expression threshold results in a list of 20 differentially expressed proteins. This might seem reasonable, but, in the worst case, all these 20 significant proteins could be among the 50 false positives from the identification step. While this is unlikely, the example illustrates that we have lost control of the FDR with respect to the hypothesis formulated above.

There are many other types of errors that can be made in a protein quantification pipeline that affect one or more of (i), (ii) and (iii). Firstly, proteins are selected for further analysis based on identification FDR. However, the identification FDR is an estimate of the evidence for the presence of proteins [33], and not a measure of how quantifiable they are i.e. their peptides being detected across conditions and being in the quantifiable range. Also, the identification FDR is often only controlled on PSM-level, which is known to underestimate the actual errors once evidence is integrated on peptide-or protein-level [10]. Secondly, missing values are rampant in data-dependent acquisition [38, 18]. Poor imputation strategies can result in unreliable results [13, 37], whereas better imputation strategies convolute the *p* value distribution from a differential expression test [13]. Thirdly, we generally rely on the fact that by averaging the peptide quantities a reliable protein quantity estimate will be obtained. However, a single misidentification, quantification error or poorly imputed value can dramatically change the results [40]. Finally, *t*-tests or ANOVAs are typically employed to search for differentially expressed proteins. In the best case, multiple hypothesis testing is applied to transform the often wrongly interpreted *p* values into more easily interpretable FDRs. It is common practice to set an a posteriori cut-off for the minimum fold change to filter out proteins with low effect size [34, 4], but this filtering actually invalidates the calculated FDRs [19]. Each of the above errors alone can cause a severe increase in false positives that remain unaccounted for and in many quantification pipelines multiple of these error sources are actually ignored. Furthermore, the overall effect of ignoring these error sources is an increase in variance leading to a drop in sensitivity of the subsequent t-tests, as mentioned above.

Bayesian methods provide a natural framework for accounting for the uncertainty at each step and propagating it to subsequent steps. For example, in the context of missing value imputation, one typically assigns a single value to replace the missing value. From this point on, this imputed value is just as reliable as any quantity that originated from an actual observation, which is intuitively ridiculous. In a Bayesian framework, we could instead assign a probability distribution over the possible values of the missing value, and when inferring the protein’s quantity we would then *marginalize* over it, that is, integrate over all the possible imputed values using their respective probabilities. This will result in a *posterior distribution*, that is *after* (post) the observation, for the protein’s quantity. This distribution will have incorporated the uncertainty due to the missing value, manifested by a larger variance than proteins without missing values.

Another important and often criticized aspect of Bayesian statistics are *prior* probabilities. Contrary to posterior distributions, prior probabilities reflect the probabilities *before* (prior) observations have been made. Prior probabilities allow us to smoothen out observations that do not fit with our initial beliefs. In the context of protein quantification, we will typically believe that most proteins are not differentially expressed. Having a single peptide exhibiting aberrant values should not immediately convince us of differential expression, as there could be a number of explanations for deviating values. However, the more peptides of that protein that show the same behavior, the more we have to override our prior belief and at some point, we will accept that the protein is differentially expressed. The controversial part of prior probabilities is that it is subjective, as each person can have a different set of prior beliefs. We can alleviate this critique by applying the empirical Bayes method, where the prior is estimated from the data.

We propose a Bayesian framework, baptized *Triqler* (TRansparent. Identification-Quantification-Linked Error Rates), formulated by a probabilistic graphical model (PGM) that combines several error models for a simple quantification pipeline, resulting in a list of significant proteins that is readily interpretable and well-calibrated.

## Methods

### Probabilistic graphical model

In a typical protein quantification pipeline (Figure 1) one starts by detecting so-called *features* in the MS1 spectra, followed by the sequence database matching of the MS2 spectra and the selection of reliable *peptide-spectrum matches* (PSMs) based on an FDR threshold. The MS1 features are then grouped by peptide identification, some type of missing value imputation is applied and the peptide quantities belonging to the same protein are then combined into this protein’s quantity Oftentimes a differential expression test is executed in the end, resulting in *p* values that subsequently should be corrected for multiple hypothesis testing, frequently followed by a fold change cutoff.

**Figure 1:**
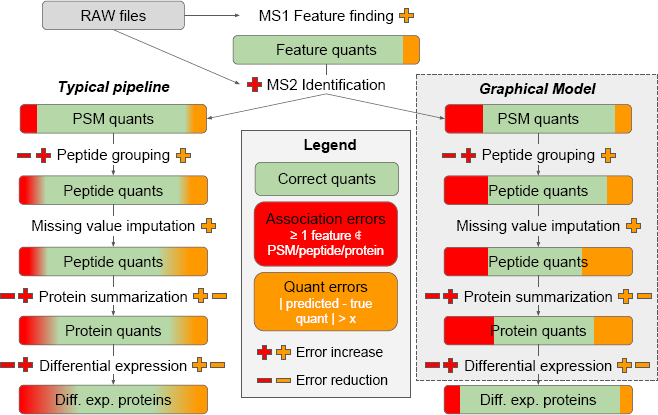
A comparison between the accounting of the errors sources of a traditional and an explicitly modeled LFQ pipeline. Each of the steps in the quantification pipeline introduces errors, which can roughly be classified as association and quantification errors. Some steps also reduce the error rate, for example by integrating evidence over multiple PSMs or peptides. In a typical pipeline, errors are implicitly propagated through different steps, resulting in a loss of control of the FDR. In contrast, the graphical model explicitly propagates uncertainty in each step and thereby controls the FDR up unto the final step. Because the uncertainty is explicitly propagated in the graphical model, we can postpone most of the filtering until the very last step. This facilitates the use of more information in each of the individual steps, without compromising the reliability of the quantifications.

In recent years, an important addition was made to the quantification pipeline in which one attempts to assign peptide identifications to features without a reliable peptide identification using similarity in retention time and precursor mass [3, 39, 1]. This greatly reduces the missing value problem but comes at the cost of having to align retention times and controlling for false discoveries. For the sake of clarity, we omit this type of inference here, but extensions to the PGM to include this are relatively simple and will be explored in future work.

We can model the quantification pipeline with a probabilistic graphical model (PGM) (Figure 2). The most intuitive way of interpreting this PGM is as a generative model for the *f*_*grn*_, which are the observed extracted ion currents (XIC):

- To assess differential expression, the main entity of interest is the mean protein quantity *z*_*g*_ of treatment group *g*.
- For each replicate *r*, we model the protein quantity, *y*_*gr*_, as coming from a distribution with mean *z*_*g*_ and variance *s*_*g*_.
- For each peptide *n*, we model the quantity, *q*_*grn*_, as the product of the ionization efficiency, *i*_*n*_, and the protein quantity, *y*_*gr*_.
- The probability that the peptide is not detected, *m*_*grn*_, depends on the peptide’s quantity, *q*_*grn*_, with lower quantities leading to higher rates of missed detections.
- The probability that the identification was correct, *t*_*grn*_, is obtained from the identification pipeline as the posterior error probability of the PSM, *c*_*grn*_.
- Finally, if the peptide was detected and correctly matched to a spectrum, *f*_*grn*_, is modelled by a distribution with mean *q*_*grn*_. If the peptide was not detected, it generates an observation *f*_*grn*_ = NaN, indicating a missing value. If the peptide was incorrectly matched, *f*_*grn*_ is generated from a distribution with the geometric mean of the observed features as mean.

**Figure 2:**
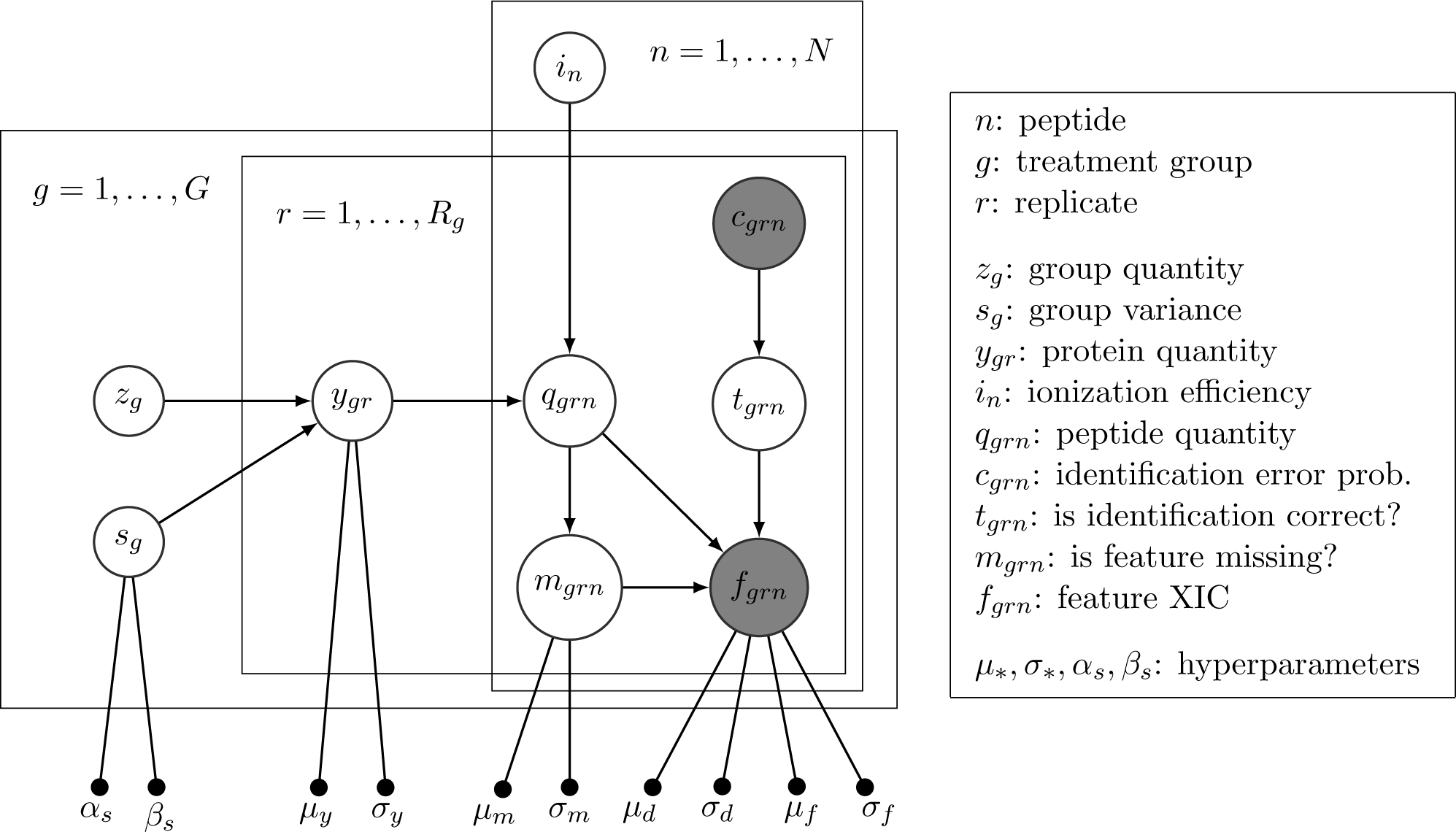
Probabilistic graphical model for the quantification of a single protein using plate notation. The protein has *N* peptides, *G* treatment groups and *R*_*g*_ replicates per treatment group. Gray nodes denote observed variables, whereas white nodes denote latent variables.

We also have 5 pairs of hyperparameters that model several underlying distributions that the random variables were drawn from. These hyperparameters were estimated from the observed data using the empirical Bayes method. These distributions were chosen based on the shape of the empirical distributions (see Supplementary Section 1). Specifically, we make use of the hyperbolic secant distribution for the feature likelihood and protein prior distributions. These distributions are similar to the normal distribution but have heavier tails, that is, they assign more probability mass to rare events. For a more detailed explanation of the variables above, the hyperparameters, the distributions employed and the hyperparameter estimation, see Supplementary Section 1.

To deal with missing values, we set a probability distribution over the imputed value using a censoring model that assigns higher probabilities to low ion currents in the event of a missing value [13, 15]. Contrary to [13], we fit a censored normal distribution of the form censnorm 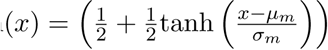.

𝒩(*x* | *μ*_*f*_,*σ*_*f*_) to the distribution of all XIC values and, furthermore, use the probability distribution directly, instead of drawing a single point estimate from the distribution. The key observation for choosing this distribution is that the empirical distribution looks like a normal distribution with mass missing at lower intensities (Supplementary Figure S2d). Note that this distribution only accounts for the “missing not at random” values, as defined in [18]. Attempts to explicitly model the “missing completely at random” values in the PGM resulted in a severe loss of sensitivity. Instead, we use the fact that the data is integrated over multiple peptides and combined with the prior on protein quantity to account for these.

Optionally, one could add prior distributions for the *z*_*g*_ and *i*_*n*_. However, in practice, it turns out that a uniform improper prior that assigns equal probability to all possible values between (–∞, ∞) works best for *z*_*g*_, which does not impose a structure on the pattern of differential expression. For the *i*_*n*_, we opt for a scheme resembling an expectation-maximization step, where we update the *i*_*n*_ with a point estimate using the geometrical average of the ratio *f*_*grn*_ and the maximum a posteriori estimates of *y*_*gr*_ (Supplementary Section 1.1). This greatly simplifies the integrals that need to be computed and works satisfactorily in practice. For a typical dataset, the execution time of Triqler is a matter of minutes, which is negligible compared to the feature extraction and peptide identification steps.

As mentioned before, *t*-tests or ANOVAs come with certain issues. We avoid these problems by testing directly for the hypothesis of interest: “What is the probability that protein *P* is correctly identified and has a fold change of at least *C*?”. We do this by combining the protein posterior error probability (PEP) of the identification step with the posterior probability for a fold change to be smaller than the threshold *C* [16]. This posterior probability can easily be calculated from the *z*_*g*_ posterior distributions, the calculation of which is outlined in Supplementary Section 1.1. Finally, we can sort the proteins by this combined PEP and calculate FDRs by simply taking the running average of the PEPs.

### Data sets

We downloaded RAW files for 3 datasets with spiked-in proteins at known concentrations, the iPRG2015 study (MassIVE ID: MSV000079843, 12 RAW files) [2], the iPRG2016 study (PRIDE project: PXD008425, 9 RAW files) [31] and a sample of the UPS1 protein mixture spiked in at 3 different concentrations in a yeast background (PRIDE project: PXD002370, 9 RAW files) [9]. We also downloaded RAW files for a clinical dataset of bladder cancer [17] (PRIDE project: PXD002170, 8 RAW files), which we will refer to as the *Latosinska* dataset.

The iPRG2015 dataset consisted of 6 known proteins of foreign origin spiked into a background of yeast at different concentrations (Table 1 in [2]). The iPRG2016 dataset featured two pools, pool *A* and pool *B* of protein fragments known as PrESTs [36], where the first sample only contained the pool *A* PrESTs, the second only the pool *B* PrESTs and the third an equimolar mixture of the two pools combined in the *A*+*B* pool. The UPS-Yeast mixture consisted of 3 samples, where a UPS1 protein mixture was spiked into a 1 *μ*g yeast background at respectively 25, 10 and 5 fmol concentration. Each of these datasets used triplicates for each sample. The Latosinska dataset consisted of 8 samples of tumor tissues of non-muscle invasive (stage pTa, *n* = 4) and muscle-invasive bladder cancer cases (stage pT2+, *n* = 4), without replicates.

**Table 1:**
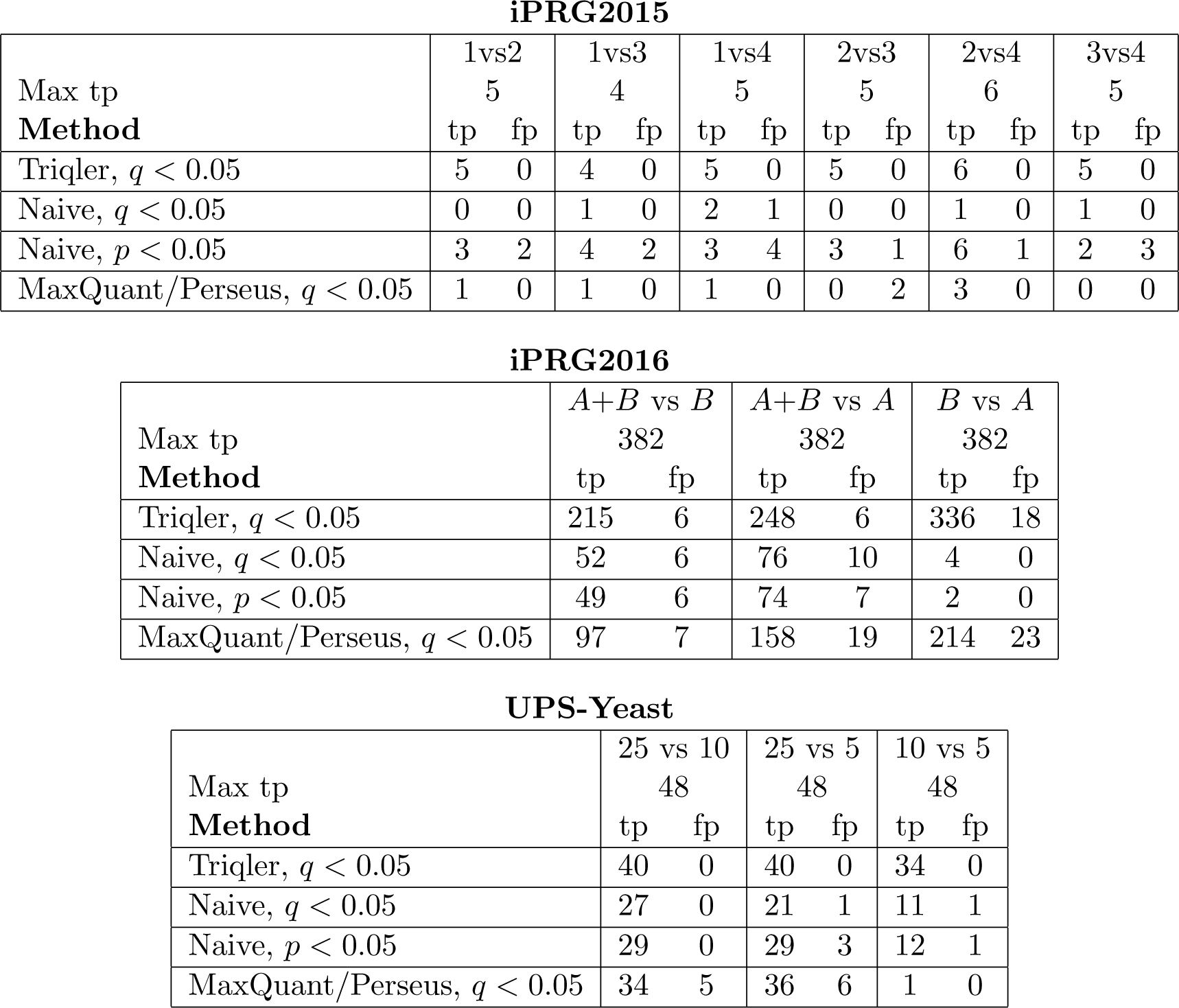
Triqler achieves a higher sensitivity than the naive method as well as MaxQuant/Perseus and controls the FDR on all datasets. Number of true and false positive significantly differentially expressed proteins at a 5% reported FDR threshold. For the naive method, an extra row was added for a *p* value < 0.05 cutoff, due to the low sensitivity at the FDR threshold for the iPRG2015 dataset.

**iPRG2015**
| Max tp<br>**Method** | 1vs2 |  | 1vs3 |  | 1vs4 |  | 2vs3 |  | 2vs4 |  | 3vs4 |  |
| --- | --- | --- | --- | --- | --- | --- | --- | --- | --- | --- | --- | --- |
|  | 5 |  | 4 |  | 5 |  | 5 |  | 6 |  | 5 |  |
|  | tp | fp | tp | fp | tp | fp | tp | fp | tp | fp | tp | fp |
| Triqler, $q < 0.05$ | 5 | 0 | 4 | 0 | 5 | 0 | 5 | 0 | 6 | 0 | 5 | 0 |
| Naive, $q < 0.05$ | 0 | 0 | 1 | 0 | 2 | 1 | 0 | 0 | 1 | 0 | 1 | 0 |
| Naive, $p < 0.05$ | 3 | 2 | 4 | 2 | 3 | 4 | 3 | 1 | 6 | 1 | 2 | 3 |
| MaxQuant/Perseus, $q < 0.05$ | 1 | 0 | 1 | 0 | 1 | 0 | 0 | 2 | 3 | 0 | 0 | 0 |

**iPRG2016**
| Max tp<br>**Method** | $A+B$ vs $B$ | | $A+B$ vs $A$ | | $B$ vs $A$ | |
| --- | --- | --- | --- | --- | --- | --- |
|  | 382 |  | 382 |  | 382 |  |
|  | tp | fp | tp | fp | tp | fp |
| Triqler, $q < 0.05$ | 215 | 6 | 248 | 6 | 336 | 18 |
| Naive, $q < 0.05$ | 52 | 6 | 76 | 10 | 4 | 0 |
| Naive, $p < 0.05$ | 49 | 6 | 74 | 7 | 2 | 0 |
| MaxQuant/Perseus, $q < 0.05$ | 97 | 7 | 158 | 19 | 214 | 23 |

Table 1: **Triqler achieves a higher sensitivity than the naive method as well as MaxQuant/Perseus and controls the FDR on all datasets.**
| Max tp<br>**Method** | 25 vs 10 |  | 25 vs 5 |  | 10 vs 5 |  |
| --- | --- | --- | --- | --- | --- | --- |
|  | 48 |  | 48 |  | 48 |  |
|  | tp | fp | tp | fp | tp | fp |
| Triqler, $q < 0.05$ | 40 | 0 | 40 | 0 | 34 | 0 |
| Naive, $q < 0.05$ | 27 | 0 | 21 | 1 | 11 | 1 |
| Naive, $p < 0.05$ | 29 | 0 | 29 | 3 | 12 | 1 |
| MaxQuant/Perseus, $q < 0.05$ | 34 | 5 | 36 | 6 | 1 | 0 |

### Data analysis

All RAW files were converted to mzML format with ProteoWizard [14]. MS1 features were detected with Dinosaur v1.1.3 [30] and assigned to MS2 spectra with an in-house python script. The iPRG2015 and iPRG2016 datasets were searched against their respective FASTA databases included in the study materials. The UPS-yeast mixture was searched against a concatenated FASTA file with the UPS1 proteins (https://www.sigmaaldrich.com/, accessed: 2018 Jan 17) and the Swiss-Prot database for yeast (http://www.uniprot.org/, accessed: 2016 Mar 15). The Latosinska set was searched against the Swiss-Prot database for human (accessed: 2015 Nov 12). The spectra were searched against their respective concatenated target-decoy database by Tide [5], through the interface of the Crux 2.1 [20] package, followed by post-processing with Percolator v3.02 [32]. All parameters in Tide and Percolator were left to their default values, except for allowing up to 2 oxidations for the iPRG2015, Latosinska and UPS-Yeast datasets.

We used the PSM-level SVM scores from Percolator as input to Triqler v0.1.2. First, the feature intensities were subjected to retention time dependent normalization, in a similar fashion to [39]. Then, several processing steps were applied inside of Triqler, ultimately resulting in a list of differentially expressed proteins. The SVM scores from Percolator were first converted to PSM-level PEPs using a Python implementation of Qvality [12], which we also used to estimate peptide-level and protein-level identification PEPs, using respectively the best SVM score per peptide as peptide score and the log of the multiplication of peptide-level PEPs as protein score [32]. For the peptide-level PEPs, we considered different charge states of the same peptide as separate peptides. Peptides with more than M missing values were filtered out and only the charge state with the best peptide-level PEP per peptide was retained. Finally, the protein-level identification PEP was combined with the posterior probability of obtaining at least a log_2_ fold change *F* and this combined PEP was then reported by Triqler.

We compared Triqler to a naive, but seemingly reasonable quantification pipeline. Here, the PSM identifications from Percolator with *q* < 0.05 were used as input and the same type of retention time dependent normalization was applied as was done before processing with Triqler. Peptides with more than *M* missing values were discarded and we only retained the charge state per peptide with the highest summed intensity. The remaining missing values were replaced by the mean of all non-missing values of the same peptide. We discarded proteins with fewer than 3 peptides and used the average of the 3 most intense peptides as the protein’s quantity. Finally, we applied a *t*-test, followed by a log_2_ fold change cutoff *F*.

We also analyzed the data with MaxQuant v1.6.1.0 [3] followed by differential expression analysis with Perseus v1.6.1.3 [35]. All parameters in MaxLFQ/Perseus were left to their default values, except that we allowed up to 2 oxidations and allowed the use of these modified peptides for quantification. Note that we did not use matches-between-runs. For the differential expression analysis, we filtered out decoy proteins and proteins with more than *M* missing values. We then applied a log_2_ transform to the intensities, imputed missing values with the default parameters and used Welch’s t-test with S_0_ = 1 (lower values of *S*_0_ typically resulted in similar or worse results).

For the iPRG2015 dataset, we allowed *M* = 5 missing values and set *F* = 0.5. For the iPRG2016 dataset, we allowed 4 missing values and used *F* = 0.8, which is just below the log_2_ fold change of 1.0 between the *A+B* pool relative to the *A* or *B* pool. For the UPS-Yeast dataset, we allowed *M* = 3 missing values and used *F* = 0.8 for the same reason as above, regarding the 5 and 10 fmol samples. For the Latosinska dataset, we allowed *M* = 4 missing values and used *F* = 1.0.

## Results

We evaluated the performance of Triqler and the naive pipeline on the four datasets described in the methods section, three controlled datasets, the iPRG2015, iPRG2016 and UPS-Yeast datasets as well as one clinical dataset, the Latosinska dataset. We also compared the results to the results of the original studies as well as those generated by MaxQuant/Perseus.

### Posterior distributions

We plotted the posterior distributions of the log_2_ fold changes between each pair of treatment groups obtained by Triqler and compared this to the Gaussian distribution obtained from the triplicate measurements for the naive pipeline. Note that, for the naive pipeline, one typically takes only a point estimate for the fold change, that is the mean of the distribution. It is, however, quite illustrative for our comparison to draw the entire distribution.

For the iPRG2015 dataset, we plotted 4 of the 6 spiked-in proteins sorted by the number of peptide identifications that were available (Figure 3). For BGAL_ECOLI and ALBU_BOVIN, which both had many identified peptides, the posterior distributions are sharp. For BGAL_ECOLI Triqler, the true fold change was in the neighborhood of the posterior distribution. Triqler seemed to underestimate the lowest concentration (sample group 1) but was at least correct about the direction of the fold change. For ALBU_BOVIN Triqler performed exceptionally well. The naive model had big troubles with the lowest concentration and tended to overestimate it, due to its missing value imputation strategy. While this conservative strategy might seem reasonable, it can lead to rather dubious results. For example, for 1vs4 for BGAL_ECOLI and 1vs2 and 2vs3 for ALBU_BOVIN it obtains the wrong sign of the fold change.

**Figure 3:**
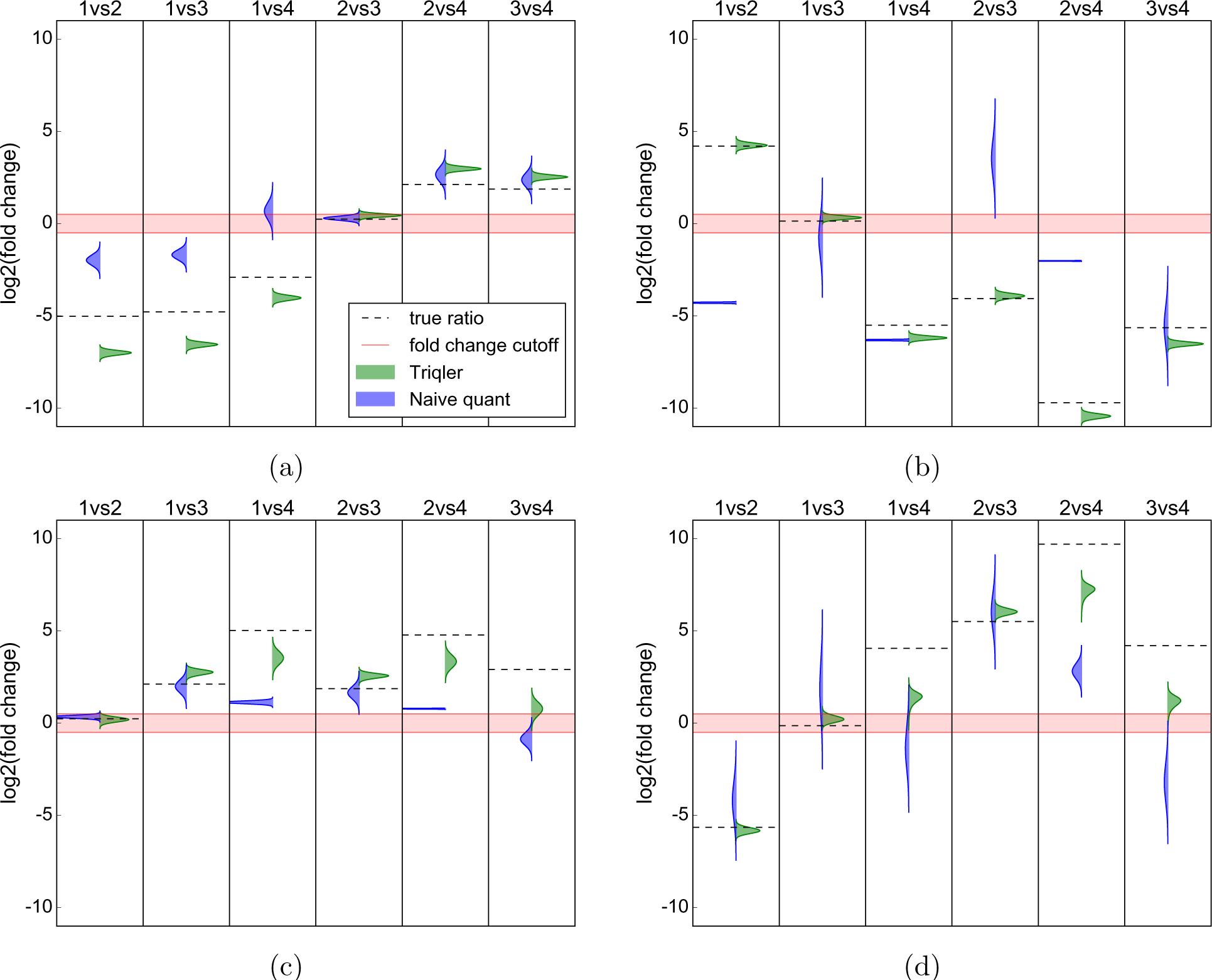
Triqler achieves reasonable estimates of the true fold changes in the iPRG2015 dataset. Posterior distributions for the fold change difference between each of the groups in the iPRG2015 dataset for 4 out of the 6 spiked-in proteins. (a) BGAL_ECOLI, with 48 identified peptides, (b) ALBU_BOVIN, with 22 identified peptides, (c) OVAL_CHICK, with 6 identified peptides, (d) CAH2_BOVIN with 4 identified peptides. The width of the posterior distribution decreases the more peptides are available.

For OVAL_CHICK and CAH2_BOVIN far fewer peptides were available for quantification. This led to broader posterior distributions for Triqler, which conforms to our intuition. In almost all cases Triqler’s posterior distribution is closer to the true fold change than the naive model. From the CAH2_BOVIN results, we can also see that the naive model will have trouble to obtain significant results when only a few peptides are available, as a *t*-test will have a hard time to separate the within-group variance from the between-group variance.

For the iPRG2016 set, the fold change of present proteins was accurately predicted (Figure 4a). We also observed a clear example of the failure of missing value imputations by the peptide’s mean abundance in the face of truly absent proteins (Figure 4b). Regardless of the amount of identified peptides, the naive method predicted values close to a fold change of 0 for these proteins. Note that Triqler predicted larger fold changes the more confident the protein identification was and moreover assigns much broader posterior distributions compared to when the protein is present in both samples. Interestingly, even when only 1 peptide identification was available, Triqler could sometimes correctly predict differential expression, although with a broad posterior distribution (Supplementary Figure S6).

**Figure 4:**
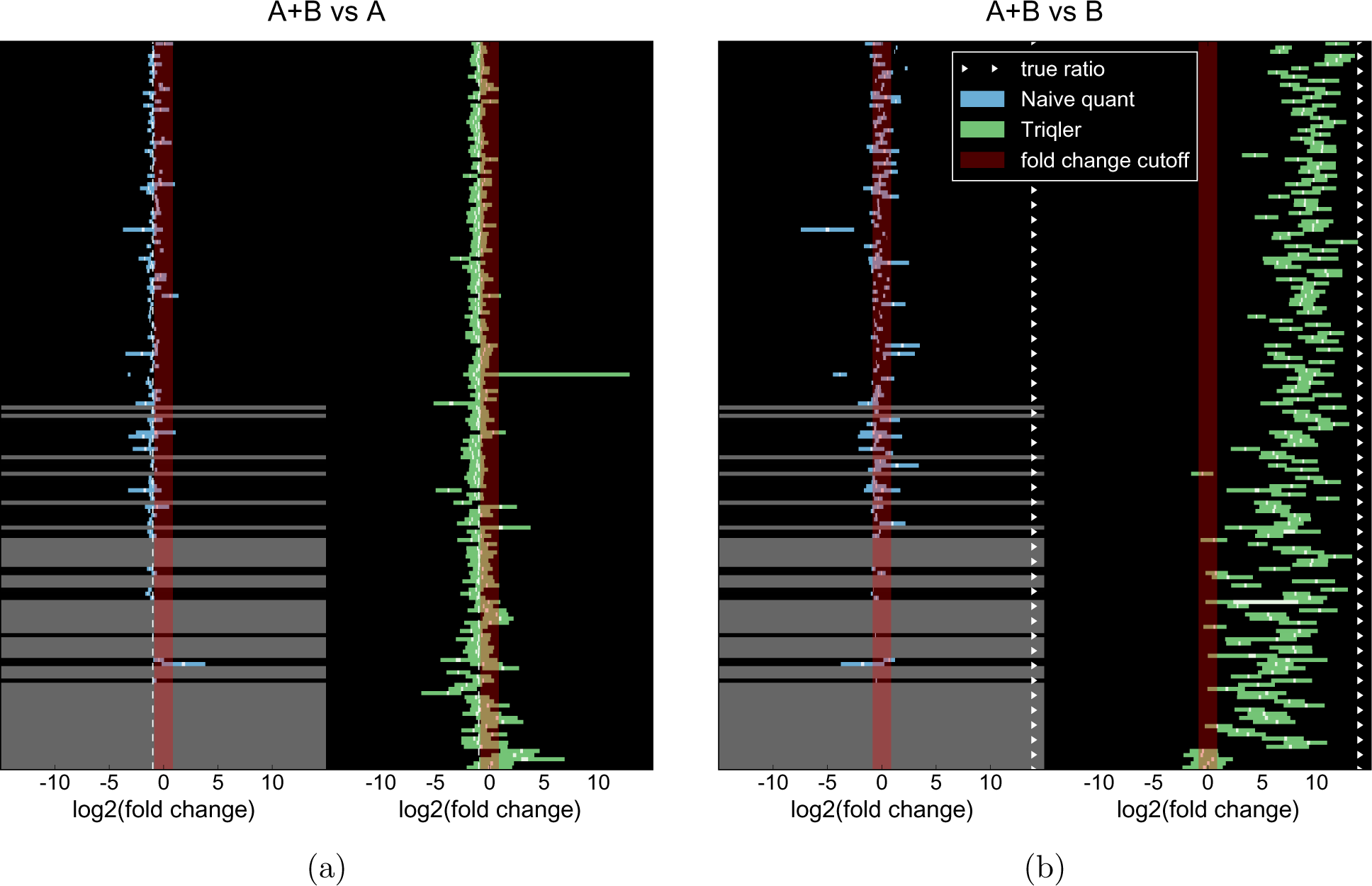
Triqler accurately estimates true fold changes (a) and predicts large fold changes for truly absent proteins (b) in the iPRG2016 dataset. Predicted fold change differences for the pool *A* spiked in proteins for (a) pools *A* + *B* vs *A* and (b) *A* + *B* vs *B*; one protein per row, sorted by protein-level FDR with the most confident protein on top. The width of the colored bars indicate the 95% confidence interval respectively credible interval [6] and the white bar inside them indicate the 10% intervals, for the naive method (left panes, blue) and Triqler (right panes, green). Grey rows for the naive method indicate proteins with fewer than 3 unique peptides. For *A* + *B* vs *A*, the true log_2_ fold change is ‐1, which the confident proteins indeed center around. The “true ratio” for *A* + *B* vs *B* only serves to indicate the sign of the expected fold change, as the protein is actually absent in pool *B*. For these cases, Triqler predicts large fold changes, whereas the naive method consistently fails to infer differential expression.

For the UPS-Yeast dataset, we again observed the broadening of the posterior distributions as the confidence in the protein identification decreased (Figure 5). We also saw that even in the case that many peptides are available, the naive model still gave rise to false negatives as can be seen for 10 vs 5 due to poor missing value imputation. Unlike for the iPRG2016 set, having only a single peptide identification was not sufficient to declare a protein differentially expressed, though in some cases there was some posterior probability covering the region of differential expression (Supplementary Figure S7).

**Figure 5:**
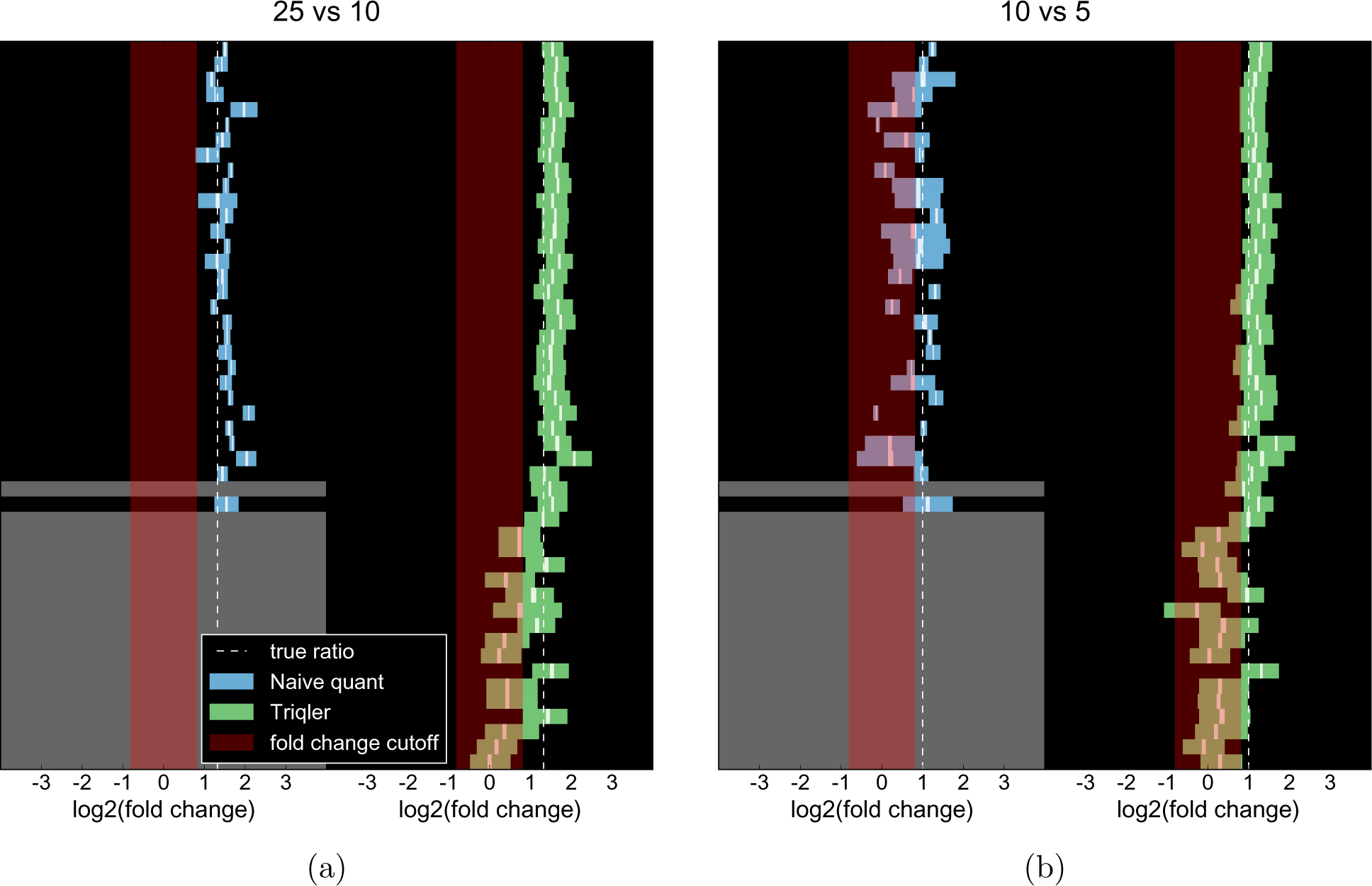
Triqler correctly predicts true fold changes for the spiked in UPS proteins for the UPS-Yeast dataset. Predicted fold changes for all 48 spiked in UPS proteins for (a) 25 vs 10 and (b) 10 vs 5; one protein per row, sorted by protein-level FDR with the most confident protein on top. The width of the colored bars indicate the 95% confidence interval respectively credible interval [6] and the white bar inside them indicate the 10% intervals, for the naive method (left panes, blue) and Triqler (right panes, green). Grey rows for the naive method indicate proteins that had fewer than 3 unique peptides. For confidently identified proteins, the posterior distributions cover the true ratios, whereas less confidently identified proteins are pulled towards the center due to the prior distribution.

### FDR control

Often, the ultimate result of a quantification pipeline is a list of differentially expressed proteins, together with an estimate of the expected proportion of false positives in this list. Unfortunately, the conventional calibration curves, which plot the observed against reported FDR, are not hugely informative. This is due to the relatively low number of truly differentially expressed proteins which gives rise to very low resolution in the low region of the FDR, where we typically set our thresholds (Supplementary Figures S8 and S9).

A more illustrative measure in these cases is the number of true positives, spiked-in proteins with the correct sign of the fold change, and false positives, spiked-in proteins with the incorrect sign and background proteins (Table 1).

We applied multiple hypothesis corrections on the *p* values coming from the naive method using Qvality. However, this approach led to very low sensitivity, and therefore we also included the results for a *p* value cutoff of 0.05, a frequently misused metric. Note that the *p* value cutoff approach actually intentionally gives up on FDR control, which could be one of the explanations behind the disproportionate amount of false positives in the iPRG2015 results.

For all 3 datasets, Triqler estimated many more true positives than either variant of the naive method as well as MaxQuant/Perseus. For the iPRG2015 dataset, Triqler obtained no false positives and no false negatives, whereas both the naive method and MaxQuant/Perseus had a much lower sensitivity. We also note that the numbers presented here for Triqler far exceed those presented by the original paper, both compared to the results from the study organizers (Figures 4 and 7 in [2]) as well as those submitted to the study by the participants (Figure 3 in [2]). This is even more remarkable when one considers that the study organizers, and likely a large proportion of the study participants as well, used some form of matches-between-runs.

For the iPRG2016 dataset, Triqler showed a reasonable estimation of the true FDR, which varied between 2.7% and 5.1%. The naive model and MaxQuant/Perseus produced too liberal FDR estimates corresponding to ≈ 10% true FDR, with much fewer true positives. The extremely low sensitivity on the *B* vs *A* comparison for the naive model was due to most proteins not making the fold change threshold filter. Furthermore, a general decrease in sensitivity could be observed due to the requirement of at least 2 (MaxQuant/Perseus) or 3 (naive method) peptides per protein. In contrast, Triqler declared several spiked-in proteins significant with 2 identified peptides and in some cases even with only a single identified peptide.

For the UPS-Yeast set, Triqler produced no false positives and was more sensitive than the naive method on all comparisons. On the other hand, we could observe that MaxQuant/Perseus had large issues controlling the FDR on this set, which was also observed in the original study (Tables 1 and 2 in the Tutorial document of the Supporting Information of [9]). The extremely low sensitivity of MaxQuant/Perseus on the UPS-Yeast set for the 10 vs 5 comparison was a consequence of the high value of *S*_0_ = 1. Setting S_0_ = 0.3 produced better results for the 10 vs 5 comparison but resulted in significantly more false positives for the 25 vs 5 comparison (Supplementary Table 3).

Note that, for both the iPRG2016 and UPS-Yeast set, there were many spiked-in proteins that did not make the 5% FDR cutoff, but still ranked above the background or entrapment proteins in terms of posterior error probability (Supplementary Figures S8 and S9). This reflects the conservative nature of Triqler due to the prior distribution which pulls estimates towards the center if not enough evidence is available.

### Analysis of the Latosinska bladder cancer dataset

The Latosinska dataset contains a comparison of muscle-invasive against non-muscle invasive bladder cancers. In the set we found 35 significant differentially expressed proteins at 5% FDR, whereas a multiple testing correction on the *p* values of the original study resulted in only a single significant protein at that FDR threshold. They did, however, find 77 proteins at a *p* value threshold of 0.05 without using a fold change cutoff. The naive pipeline did not find any significant proteins at the FDR cutoff, and only found 10 proteins with a *p* value below 0.05 and a fold change cutoff of 1.0. To assess the soundness of these significant proteins, we analyzed the concerning proteins with the functional annotation chart tool from DAVID 6.8 [11]. For Triqler, we used the 35 significant proteins below 5% FDR, for the original study, we used the 77 significant proteins below the *p* value cutoff of 0.05. Each of these lists was searched against the respective background of identified proteins and the categories selected in DAVID by default. A 5% term-level FDR threshold (p values corrected with the Benjamini-Hochberg correction) was applied to assess the significance of terms.

The 77 proteins of the original study showed no enriched terms, with the most significant term coming in at 30% term-level FDR. In contrast, the 35 significant proteins from Triqler resulted in 5 significant terms (Supplementary Table S1). Using higher FDR thresholds for the calling of significant proteins of 10% (58 proteins) and 20% protein-level FDR (115 proteins) resulted in 4 and 17 significant terms respectively (Supplementary Table S2 and S3). Moreover, analysis with MaxQuant+Perseus resulted in 4, 11 and 15 significant proteins at 5%, 10% or 20% protein-level FDR threshold respectively, with all but one of these significant proteins also identified at the 5% protein-level FDR threshold by Triqler. No significant terms could be found for the 5% and 10% protein-level FDR thresholds, and only 1 significant term at the 20% protein-level FDR threshold (Supplementary Table S4).

## Discussion

We have presented Triqler, a Bayesian model for protein quantification and differential expression analysis. The model takes information on different sources of errors into account and propagates it all the way up to the final list of differentially expressed proteins. Specifically, the model solves the problem of having separate identification and differential expression error rates and presents a unified error rate that covers both. It avoids common pitfalls of quantification pipelines and introduces the concept of posterior probabilities as a replacement for the statistically unsound fold change cutoff. Furthermore, contrary to many Bayesian models, the execution of our pipeline only takes a matter of minutes.

Specifically, our model integrates out missing values instead of imputing point estimates. This approach facilitates the quantification of proteins that are absent, such as in the iPRG2016 dataset, or present in low concentrations, such as in the iPRG2015 dataset. At the same time, it avoids false positives that typically arise due to poor missing value imputation methods, for example by imputation by the limit of detection. Furthermore, the use of empirical Bayes allows data to speak for itself through the prior distributions, rather than setting hard thresholds based on heuristics. This, for example, allows proteins with only a single identified peptide to be informative enough to be considered for differential expression in some experiments, whereas they will be sent straight to the trash bin in others.

However, some care does have to be taken with fitting distributions to the data. For example, the censoring distribution has 4 free parameters and might be sensitive to overfitting. We can currently not ascertain the correctness of these distributions but the results so far have been encouraging. For future research, it would be interesting to compare our approach with other Empirical Bayesian approaches for missing values, such as [15]. We have also observed problems when processing an engineered dataset in which lysates from different organisms were mixed at different concentrations in each condition [40]. The lack of a background that remained constant caused problems for the hyperparameter estimation of the censoring distribution and also pulled the expressions of proteins with few peptides to the mean due to the prior. On the other hand, the model was able to handle the iPRG2016 dataset which also lacked a constant background, as only the spiked-in proteins were present in the sequence database. More experiments are needed to test the robustness of the method, for example, on smaller datasets and datasets with different fractions of a constant background.

A nice feature of PGMs is that they generally are robust to the selection of parameter distributions, as the model is integrated over the parameters’ distributions. However, we estimated the hyperparameters with Empirical Bayes, instead of employing the usual practice of having extra hierarchical layers with distributions from which the hyperparameters themselves are drawn from. This would have complicated the computation such that numerical integration would no longer be feasible, but might be the reason why the model is not robust to exchanging the distributions for other ones. Replacing the hyperbolic secant distributions with normal distributions resulted in more false positives due to their light tails that assigned too low probabilities to rare events, specifically, for the feature likelihood. Conversely, using the Cauchy distribution, a heavier-tailed distribution, also produced more false positives. This was caused by having too weak protein priors that were easily overturned by missing values. Comparing the empirical distributions with the distribution fits does show that the tails are too light and too heavy for the normal and Cauchy distribution respectively.

Another point of caution is the choice of the log_2_ fold change cutoff. If one sets this too low, the posterior distributions could be of comparable or even larger width than the region of non-significance, causing some reported probability of differential expression even when the distribution is practically centered around zero as can be seen for the low confident proteins in the bottom rows of Figure 4. To avoid this, one should verify that the threshold is above 2 – 3 times the average standard deviation of the posterior fold change distributions. Additionally, one could filter out proteins individually that have too large of a standard deviation, though we have refrained from doing so here.

The presented comparison against a naive pipeline is by no means meant as a benchmark, but rather as an illustration of how seemingly reasonable choices can lead to very poorly calibrated results with low sensitivity. There are many algorithms and methods available that undoubtedly would result in better performance than the naive method presented here. For example, there are more advanced missing value imputation methods [13], protein summarization techniques [40, 41] and statistical tests [19, 25]. Each of these algorithms solves parts of the problems of protein quantification, but, aside from potential individual shortcomings, the need to combine them with other methods, down-and upstream, will almost inevitably lead to a loss of control of the FDR.

The graphical model has the benefit of explicitly modeling sources of error, which makes it easier to identify underlying assumptions and extend the model with new error sources. One particular source of error that is currently left out is the possibility of a feature to be incorrectly matched to a spectrum, which could, for example, be added by an extra node into *t*_*grn*_, the binary variable that indicates whether the feature came from a random peptide. One could also envisage extensions of the model to incorporate, among others, shared peptides [8, 29], matches-between-runs [3, 39, 1] and data independent acquisition data [26].

The posterior distributions have the ability to make the uncertainty in fold change explicit, rather than having only a point estimate that might hide a very large uncertainty. They have the added benefit that they conform to our intuition regarding probabilities in contrast to, for example, *p* values. These distributions can be summarized into a single posterior probability of obtaining a certain fold change, but they could also be fed into downstream applications, such as pathway analysis or development of biomarker assays while retaining the information regarding their uncertainty. The functional annotation analysis at 20% FDR threshold on the Latosinska dataset highlights this potential of propagating information below arbitrary thresholds, which would normally be discarded.

## Acknowledgements

We would like to thank Jonathon O’Brien, Harvard Medical School, and Andrew Roth, University of Oxford, for thoughtful discussions on Bayesian statistics. L.K. was supported by a grant from the Swedish Research Council (grant 2017-04030).

